# Brain structure-function relationships across the human lifespan based on network eigenmodes

**DOI:** 10.1101/2023.03.08.531719

**Authors:** Yaqian Yang, Shaoting Tang, Xin Wang, Yi Zhen, Yi Zheng, Hongwei Zheng, Longzhao Liu, Zhiming Zheng

## Abstract

While brain function is considered to be tightly supported by the underlying structure, the connectome-based link estimated by current models is relatively moderate, leaving the structure-function relationship an ongoing challenge in neuroscience. Here, by proposing a novel mapping method based on network eigendecomposition, we present a concise and strong correspondence between structure and function. We show that the explanation of functional connectivity can be significantly improved by incorporating interactions between different structural eigenmodes, highlighting the potential importance of collective, higher-order coupling patterns between structure and function. We also demonstrate the pronounced advantage of the present mapping in capturing individual-specific information, and apply it to assess individual differences of structure-function coupling across the lifespan. We find that structure-function liberality weakens with age, which is driven by the decreases in functional components that are less constrained by anatomy, while the magnitude of structure-aligned components is preserved. Our results contribute to a more refined understanding of structure-function coupling and how it evolves with age.

## Introduction

The human structural connectome promotes communication among distributed cortical regions, giving rise to richly patterned neural synchrony that is thought to support a wide range of cognitive functions and behaviors (*1, 2*). Characterizing the relationship between brain structure and function is a fundamental question in neuroscience, which is instrumental for understanding how cognitive processes emerge from the underlying anatomical pathways and for advancing the treatments for neurological and psychiatric diseases (*3*). With the development of network science and imaging techniques, brain structure-function relationships are increasingly investigated using macroscale structural connectivity (SC) and functional connectivity (FC) networks, which characterize the physical pathways and temporal synchrony between brain regions, respectively (*4*). A number of studies (*5–7*) have shown that there exists a significant correlation between these two measures, where SC appears to act as a skeleton that constrains FC.

Multiple models have been proposed to explore how the FC network is coupled with the SC network, ranging from the simplest one-to-one mapping (*8*) using statistical correlations to more sophisticated biophysical models (*9, 10*) that derive functional connectivity from large-scale simulations of neural activity dynamics. Communication models (*11, 12*) fall between these two extremes, where functional connectivity is conceptualized as a weighted superposition of communication events over the structural network, with the forms of communication ranging from the shortest path routing (centralized) to signal diffusion (decentralized) (*13*). This approach achieves higher accuracy than the direct correlation method and lower complexity than the biophysical models, and as a result, becomes increasingly common in SC-FC mapping studies. Besides, another appealing tool for SC-FC mapping is the eigenmode approach (*14, 15*). This approach exploits a simple linear model that represents the FC network as a weighted combination of structural eigenmodes but achieves a high prediction accuracy comparable to sophisticated nonlinear models. These eigenmodes summarize structural connectivity into frequency-specific spatial patterns, opening a new avenue to explore structure-function relationship by decomposing functional signals into the eigenspectrum of the structural connectome (*16, 17*).

In addition to modeling advances, structure-function relationships have also been applied to investigate the effects of cognitive tasks (*18, 19*), lesions (*20, 21*), neurological disease (*22, 23*), development and aging (*24–26*). As one of the main goals of SC-FC mapping models is to capture the essential principle of how structure and function are related, a natural expectation is that the estimated structure-function relationships would have behavioral significance and could reflect the effects of manipulations and perturbations (*27*). Indeed, some recent studies have revealed associations between SC-FC correlations and various cognitive traits. One such study shows that increased alignment between structure and function is related to better cognitive flexibility (*18*). Other studies suggest that weaker SC-FC coupling is related to increasing awareness levels (*23*) and better recovery after severe brain injury (*21, 28*). Moreover, the strength of structure-function coupling is demonstrated to be heritable and to vary with subjects’ sex and age (*24–26, 29*).

Although SC-FC mapping has been fruitfully investigated and widely applied, the current literature is subject to the relatively moderate correspondence between brain structure and function. SC rarely explains more than 50% of the variance in empirical FC (*27*), which implies that, to a great extent, the mechanisms underlying the formation of functional connectivity remain elusive. In particular, recent work unifying various eigenmode approaches reports that none of the individual-specific SC-FC mappings could outperform a reference mapping that does not utilize structural information but just returns the group-average FC (*14*), raising important concerns that individual variance in structure-function relationships may not be accurately quantified. Another study comparing a large number of communication models shows that whole-brain FC is poorly predicted from structure in individuals, irrespective of predictors (*26*). As strong alignment between predicted and empirical functional networks appears desirable to ensure the fidelity of the captured information, this modest explanatory power is unfavorable for the refined investigation of structure-function relationships and for further applications to individual differences associated with behavior and cognition.

Why this imperfect link between SC and FC? There exist two intriguing hypotheses. The first one is that SC and FC may be indeed decoupled to some extent, implying that function cannot be completely predicted by structure alone. Several studies on regional structure-function relationships have shown that structure and function are tightly coupled in primary sensorimotor cortex but decoupled in polysensory association cortex (*30, 31*). This gradual divergence closely follows representational and cytoarchitectonic hierarchies, in parallel to a functional gradient (*32*) that associated cortical organization with a spectrum of increasingly abstract cognitive functions, raising a possibility that the observed structure-function divergence may be a fundamental property of the brain organization. The alternative hypothesis is that SC and FC may be tightly coupled but current models leave out information requisite for precise prediction. Multiple studies have revealed the important roles of microstructural properties in functional interactions (*33–35*), and the explanation of function is improved by incorporating information on gene co-expression (*36*), raising the possibility that SC-FC correspondence could be enhanced by more nuanced models that encompass biological details. Indeed, recent studies (*37, 38*), using the machine learning approach and high-frequency eigenmodes, have achieved substantially higher structure-function prediction accuracy than previously suggested. Nevertheless, high accuracies of these approaches often come with high execution time and model complexity, and the essential principles of the FC organization are still unclear.

Therefore, whether, and if so, how to establish a simpler and tighter link between SC and FC remain an important unsolved issue in the investigation of structure-function relationships. Here, we attempt to shed light on this question with a novel mapping framework that interprets the essential pattern of functional interactions in the context of the structural eigenspectrum. Different from the previous SC-FC mappings that keep the interregional connectivity central, our approach concentrates on the inherent patterns of brain functional interactions, which not only effectively reduces the complexity of the mapping procedure but also may yield improved robustness against weak spurious connections induced by noise (*39*). In this way, we aim to provide a more concise and accurate quantification of brain structure-function relationships and show how SC-FC coupling changes over the human lifespan by applying the captured individual-specific information.

We first show that functional brain interaction patterns are dominated by a few functional eigenmodes, and construct a link between structure and function by projecting the most contributing functional mode into the eigenspectrum of the SC network. The mapping procedure, while simple and feasible easily, achieves significantly higher accuracy than the conventional eigenmode approach and communication model, implying that structure-function coupling appears to be substantially tighter than previously suggested. Intriguingly, and reinforcing the idea of regional heterogeneity in structure-function relationships, we also observe system-specific effects in the predictability of FC in all three approaches, with primary sensory regions overall exhibiting higher prediction accuracies than polysensory association regions. Next, to examine whether the present mapping is able to capture additional information not explained by the mean, we benchmark it against a reference mapping that just returns the group-average FC network. We find that the proposed mapping yields prediction accuracy comparable to the mean mapping in an age-homogenous population (28.8 *±* 9.1 years) while achieving improved accuracies in a population across the lifespan (35 *±* 20 years). In contrast to that, the eigenmode and communication models exhibit significantly worse prediction performance than the reference mapping in both populations. These findings demonstrate the unique capability of the proposed mapping to capture individual-specific information on functional connectivity, which holds great potential for a deeper understanding of individual variability in structure-function relationships. Finally, we apply the proposed SC-FC mapping to explore the evolving properties of structure-function relationships. We find that functional portions that deviate from structural connections decrease with age while the structure-aligned portions are preserved, manifesting as gradually weakened structure-function liberality across the human lifespan.

## Results

Our analyses are organized as follows. We begin by presenting a simple and accurate SC-FC mapping, analyzing both whole-brain and regional SC-FC coupling, and comparing the present mapping to the eigenmode approach and communication model (the first two sections). We then examine additional information not captured by the mean in terms of whether the subject-specific SC-FC mapping outperforms the mean FC mapping (the third section). Finally, we analyze how SC-FC relationships evolve across the human lifespan (the fourth section). We employ two independent datasets in this study. The first one comes from the Department of Radiology, University Hospital Center and University of Lausanne (LAU), including structural and functional data from 70 healthy young participants (28.8 *±* 9.1 years). We quantify whole-brain and regional SC-FC coupling, verify the performance of the proposed method and quantify differences between different types of SC-FC mappings in this dataset. The second one is the Nathan Kline Institute (NKI)/Rockland Sample public dataset, which includes 196 healthy participants aged from 4 to 85 years. We exploit it to confirm the method’s reproducibility and explore inter-individual variation in SC-FC relationships across the human lifespan. For details of data processing and network reconstruction, see Materials and Methods.

### SC-FC mapping through structural and functional modes

Accurate quantification of SC-FC relationships is imperative for understanding how interacting brain circuits support cognitive functions and how structure-function relationships change across developmental and aging stages. However, the correspondence between structure and function is modest in current models, limiting the mechanistic insight into the tethering between the organization of physical connections and the pattern of functional interactions.

Here, we characterize a simple but strong structure-function relationship via the graph spectra of SC and FC networks. As illustrated in Fig. 1, the eigendecomposition of the SC network provides a set of eigenvectors sorted in decreasing order of their eigenvalues, representing distinct inherent modes of the structural connectome. The alignment of these eigenmodes with the anatomical connections is measured by their eigenvalues (*18*). Typically, the eigenvalue will be positive if the corresponding structural mode is strictly constrained by the underlying structural connectivity and negative if the structural mode is misaligned with anatomy (Fig. 1a). We employ these mutually orthogonal eigenvectors as a parsimonious basis for the brain FC network, which is also decomposed into its constituent eigenmodes with eigenvalues reflecting the contribution of each mode (*40*). Larger eigenvalues indicate greater contributions of functional modes to the FC network. As shown in Fig.1b and Supplementary Fig. S1, we find that the contribution of functional modes to the formation of complex functional interactions is extremely heterogeneous (Fig.1b), with just a few functional modes almost explaining the whole brain functional connectivity (Supplementary Fig. S1). Accordingly, we project the functional mode with the largest contribution into the eigenspectrum constituted by structural modes, providing a simple mapping procedure to transform the SC network into the richly patterned functional network. The predictors are structural eigenmodes. The observation is the whole FC network that is approximated by the most contributing functional mode. Parameters can be easily computed in closed form, see Materials and Methods for more details. Note that though we focused on the largest functional mode in this paper, future work could naturally incorporate more functional modes in SC-FC mapping and tune the balance of prediction accuracy and computational complexity according to the tasks (Supplementary Fig. S2).

**Fig. 1.**
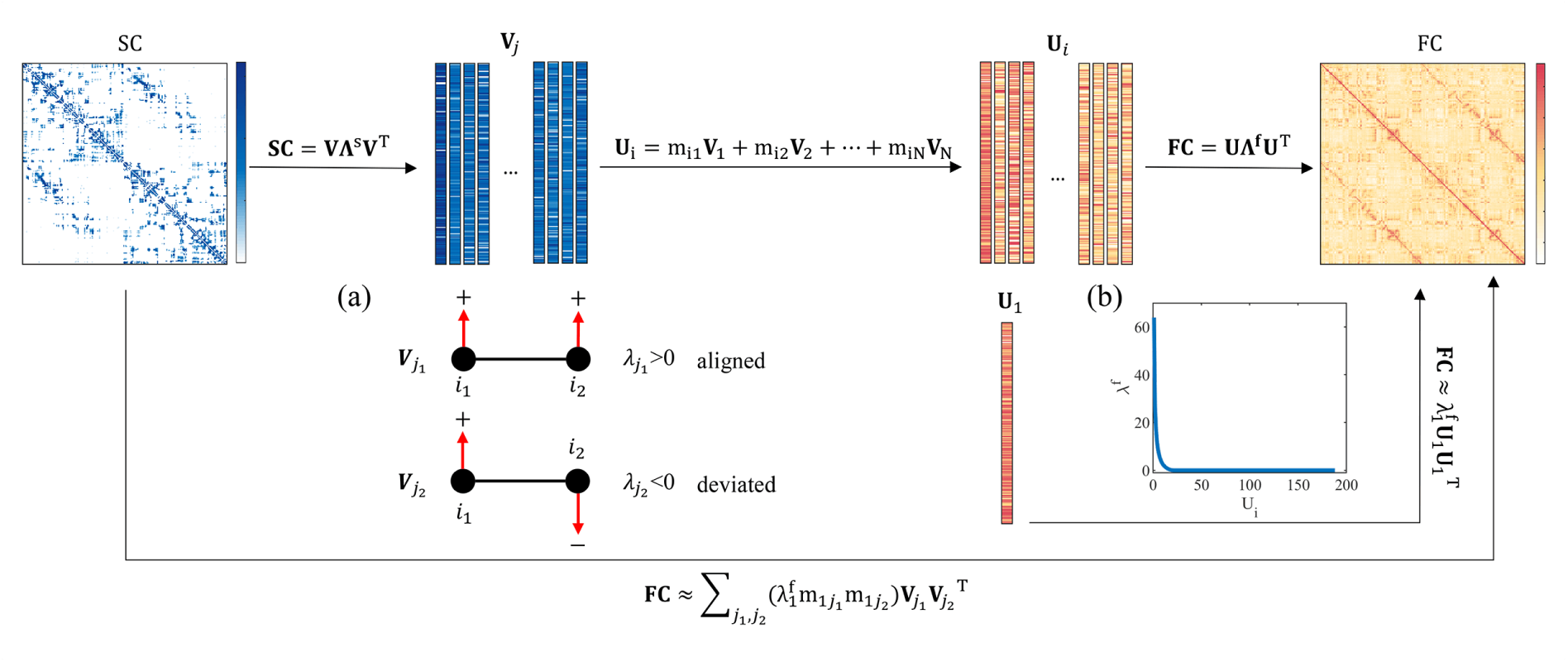
Method pipeline. Through the eigendecomposition of the structural network, we obtain a series of eigenmodes sorted in decreasing order of eigenvalues. The magnitudes of eigenvalues indicate the degree to which structural modes align with the underlying anatomy, with the large positive values corresponding to strict alignment and negative values corresponding to the deviation from SC (**a**). These mutually orthogonal structural modes can be considered as a parsimonious basis for the FC network which is also decomposed into its constituent eigen-modes accompanied with eigenvalues reflecting their contributions (**b**). The structure-function mapping is constructed by projecting the most contributing functional mode into the structural eigenspectrum.

### Enhanced explanation of the FC network

To assess the performance of the present method, we construct whole-brain SC-FC mapping for each subject from the LAU dataset and compare its performance with the conventional eigen-mode approach that utilizes the same predictors, i.e., structural eigenmodes. The Pearson correlation coefficient R between the predicted and empirical FC networks is calculated to evaluate the prediction performance and the strength of structure-function coupling. As shown in Fig. 2a, we find that the proposed method yields significantly higher prediction accuracy relative to the eigenmode approach (proposed: *R* = 0.59 *±* 0.09, conventional: *R* = 0.21 *±* 0.03, paired t-test *p <* 10*^−^*^10^), demonstrating a substantially stronger structure-function correspondence than previously implied. Note that, while the number of parameters is the same in these two techniques, the proposed SC-FC mapping could incorporate more information by introducing the cross terms of the structural modes. Thus, these findings also suggest an important role of the interaction between different structural modes in the FC formation, emphasizing the collective, high-order relationship between structure and function that transcends a linear superposition of structural eigenmodes.

**Fig. 2.**
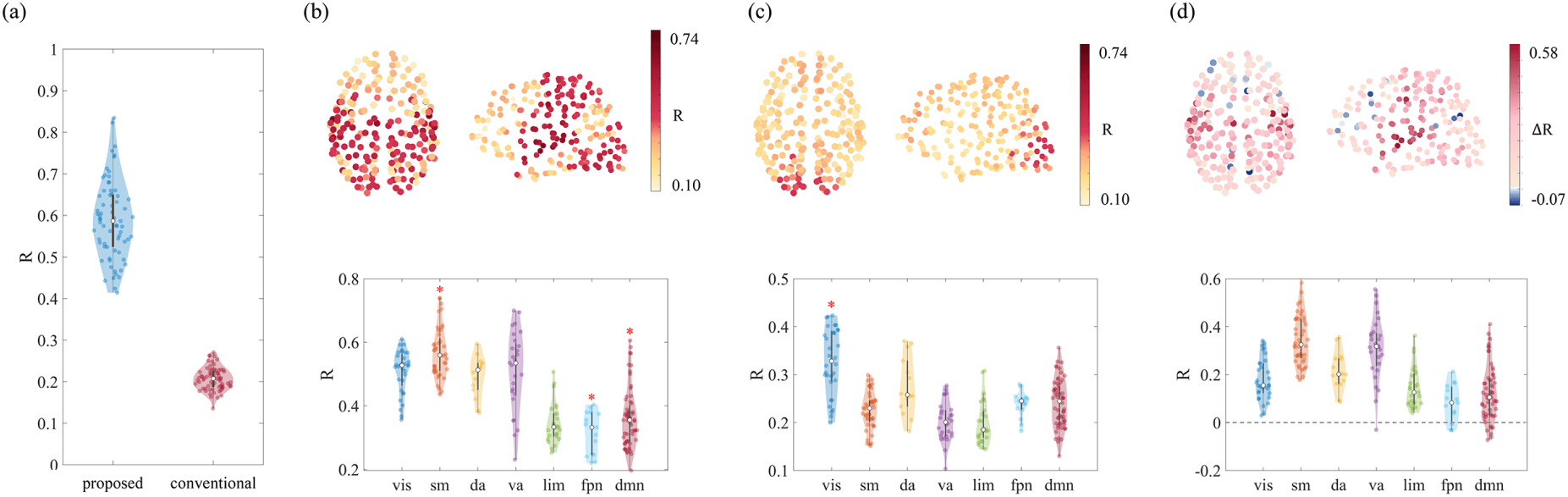
Whole-brain and regional performance of the proposed mapping. (**a**) The prediction accuracy of the proposed method vs. the conventional eigenmode approach for all individual subjects. In each violin plot, the box indicates the interquartile range and the empty circle indicates the median value. (**b**) The spatial pattern of SC-FC coupling estimated by the proposed method (upper panel) as well as the distribution of R over regions aggregated by seven resting-state networks (RSNs) proposed by Yeo et al (lower panel; asterisks indicate statistical significance). (**c**) The spatial pattern of SC-FC coupling estimated by the conventional eigen-mode approach (upper panel) as well as the distribution of R over regions aggregated by seven RSNs (lower panel). (**d**) Regional differences between prediction accuracies of the proposed and eigenmode methods. Seven RSNs include visual (vis), somatomotor (sm), dorsal attention (da), ventral attention (va), limbic (lim), frontoparietal (fpn), default mode (dmn) networks.

We then consider the strength of regional SC-FC coupling, which is estimated as the correlation between the same region’s predicted and empirical FC profiles. As shown in Fig. 2b and c, we find that regional structure-function coupling varies considerably across the cortex in both mapping approaches. In the proposed SC-FC mapping, regions with high prediction accuracy are concentrated in visual cortex, supertemporal cortex, and somatomotor (preccentral and postccentral) cortices, whereas regions in precuneus, cingulate, and prefrontal cortices exhibit relatively low prediction accuracy. To further characterize these findings at the level of functional systems, we aggregate accuracy R by seven resting-state networks proposed by Yeo et al (*41*) and compare the network-specific mean R to those generated by spatially-constrained permutation (spin test) (*42*). We find that FC profiles of regions in the somatomotor network are better explained than the null distribution while FC profiles of regions in the frontoparietal and default mode networks are worse explained than explained by chance (10,000 permutations; all FDR-corrected *p <* 0.01). Such system-specific effects also appear in SC-FC coupling estimated by the conventional eigenmode approach, with FC profiles of regions in the visual network significantly better explained than those of other regions (10,000 permutations; FDR-corrected *p <* 0.01). We further compare the regional R estimated by the proposed mapping with that estimated by the previous eigenmode approach (Fig. 2d). We find that, while the structural predictors and model complexity are identical in these two techniques, the proposed method generally outperforms the eigenmode approach, with the FC profiles of 76 *±* 8% of regions being better explained by introducing the interactions between different structural modes.

Next and for completeness, we compare the proposed mapping with a communication model that incorporates a large number of predictors characterizing the geometric, topological, and dynamic relationships between regions (Fig. 3a). Broadly, these structurally informed predictors include flow graphs (parameterized at different timescales) (*43*), mean first passage times (*44*), communicability (*45, 46*), matching index (*47*), path transitivity (parameterized at weight-to-cost transformations) (*12*), search information (parameterized at weight-to-cost transformations) (*48*), Euclidean distance, and the shortest path length. As shown in Fig. 3b, we find that the prediction accuracy of the proposed method is significantly higher than that of the communication model (communication: *R* = 0.30 *±* 0.04, paired t-test *p <* 10*^−^*^10^), confirming the strong explanatory power of the present SC-FC mapping. We also observe regional heterogeneity in structure-function coupling estimated by the communication model, with regions in the visual network exhibiting higher prediction accuracy than regions in other functional systems (10,000 spatially-constrained permutations, FDR-corrected *p <* 0.01; Fig. 3c). Furthermore, we show that the present method yields higher prediction accuracy than the communication model across a wide range of cortex, with FC profiles of 67 *±* 10% of brain regions being better explained by the proposed mapping (Fig. 3d).

**Fig. 3.**
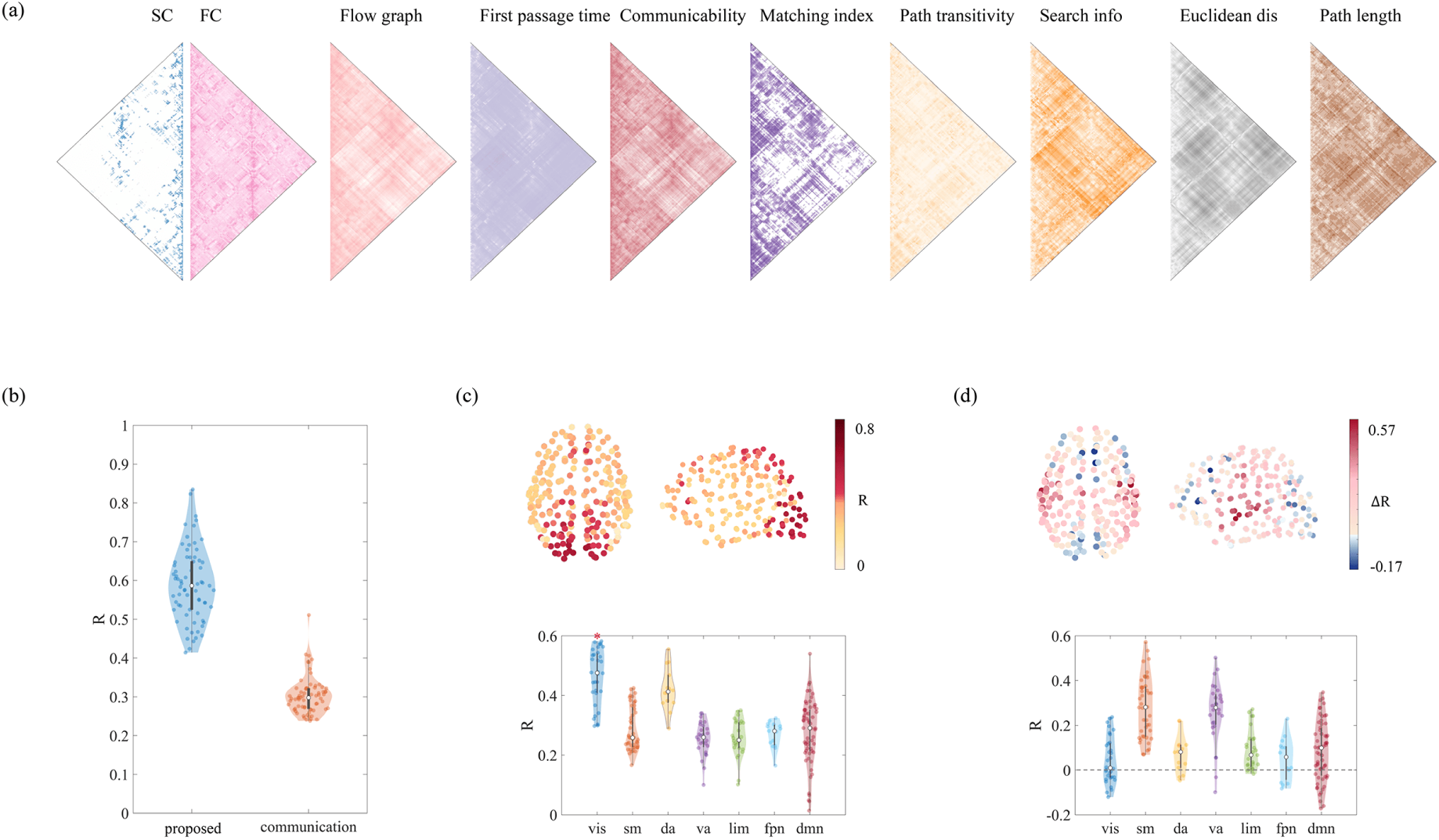
Comparison with the communication model. (**a**) A communication model that incorporates a large number of structurally informed predictors characterizing the geometric, topological, and dynamic relationships between regions. (**b**) The prediction accuracy of the proposed method vs. the communication model for all individual subjects. In each violin plot, the box indicates the interquartile range and the empty circle indicates the median value. (**c**) The spatial pattern of SC-FC coupling estimated by the communication model (upper panel) as well as the distribution of R over regions aggregated by seven RSNs (lower panel; asterisks indicate statistical significance). (**d**) Regional differences between prediction accuracies of the proposed and communication methods.

Collectively, these results confirm the validity of the proposed SC-FC mapping, providing new evidence that brain structure-function correspondence is substantially stronger than previously implied. Interestingly, regional heterogeneity is preserved in this enhanced structure-function relationship, closely aligned with the previous findings (*30, 31*) that primary sensory regions overall exhibit tighter SC-FC coupling than polysensory association regions. Further more, as the present SC-FC mapping is constructed via the largest functional mode, our findings also suggest the powerful role of functional modes as a highly effective tool for the dimensionality reduction of FC networks. The reproducibility of findings shown above are verified in an independently collected dataset (NKI; Supplementary Fig. S3 and S4).

### Unique advantage in capturing individual-specific information

Having confirmed the effectiveness of the proposed method, we next seek to assess whether it is able to characterize inter-individual variation in functional connectivity. To this end, we introduce a reference mapping that simply returns the group-average FC network (Fig. 4a). This mapping neither utilizes structural information nor preserves inter-individual variation, thereby providing a benchmark against which the relative performance of individual-specific structure-function mapping could be measured. The correlation coefficient R between the mean FC and each individual’s empirical FC is computed, generating a null distribution for the prediction accuracy of individual-specific mappings that utilize structural information. The accuracy R of the proposed method, the eigenmode approach, and the communication model are then compared to this null distribution via the paired t-test, with the t-statistic reflecting the significance of the difference.

**Fig. 4.**
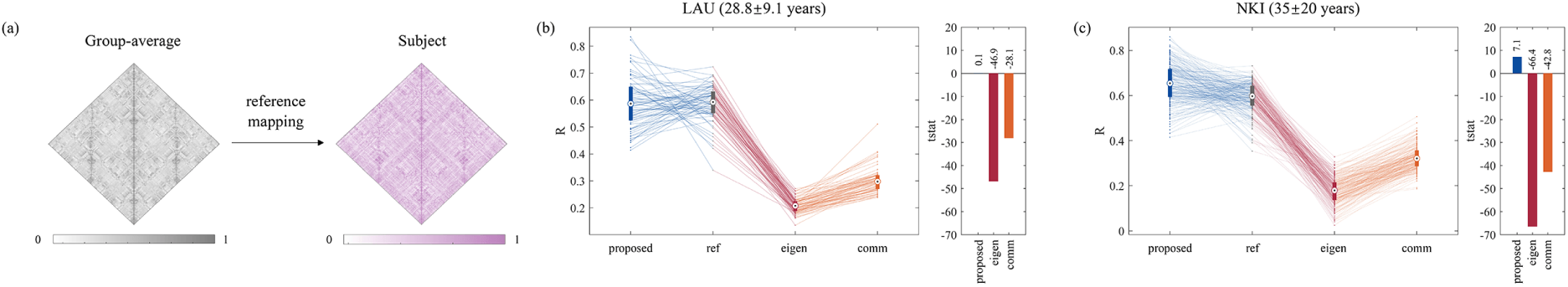
Individual-specific mapping between structural and functional networks. (**a**) A reference mapping that only returns the group-average FC network. (**b**) Comparison of different mapping methods in the LAU dataset. Left panel: Prediction accuracies of individuals’ FC networks through the proposed, reference, eigenmode, and communication methods. In each boxplot, the box indicates the interquartile range and the empty circle indicates the median value. Paired points in distinct boxplots connected by lines represent the same individual. Right panel: Prediction accuracies of the three personalized mappings are benchmarked against the reference mapping. (**c**) Comparison of different mapping methods in the NKI dataset.

We employ two independent datasets to perform the comparison. The first one is the LAU dataset, which consists of a homogeneous population of roughly the same age range (28.8 *±* 9.1 years). The second one is the NKI dataset, which comprises a relatively heterogeneous population across the human lifespan (35 *±* 20 years). Interestingly, we find that not all SC-FC mappings that utilize subject-specific structural information and preserve inter-individual variation outperform the mean mapping (Fig. 4b and c). Instead, in a homogeneous population, the mean mapping appears to serve as an apparent glass ceiling for FC prediction, with the accuracy R of both the eigenmode approach and the communication model significantly lower than that of the reference mapping (reference: *R* = 0.58 *±* 0.06; paired t-test *P <* 10*^−^*^10^). One possible explanation may be that, in these mapping procedures, individual-specific information introduced by structurally informed predictors may not be sufficient to overcome the noise introduced by high susceptibility artifacts in structural connectomes. Notably, the proposed SC-FC mapping still performs well, achieving comparable prediction accuracies to the mean mapping (paired t-test P=0.90). Furthermore, in a heterogeneous population, the prediction accuracy of the present mapping is higher than that of the reference mapping whereas the eigenmode and communication methods still fail to outperform the reference mapping (all paired t-test *P <* 10*^−^*^10^), indicating the unique capability of the proposed method to capture additional subject-specific information not explained by the mean.

Collectively, these results suggest that individual-specific SC-FC mapping appears to be quite susceptible to noise in structural imaging data, resulting in a significant reduction in prediction accuracies of previous mappings (eg., the eigenmode and communication approaches) compared to that of the reference mapping. Nevertheless, the proposed SC-FC mapping overcomes this potential pitfall to a great extent and even captures additional individual-specific information in a less homogeneous population, providing more accurate quantification of structure-function relationships at an individual level.

### Weakened structure-function liberality across the human lifespan

The present mapping has shown pronounced advantages in characterizing individual-specific information, which raises the prospect of a more nuanced investigation of individual differences in structure-function coupling. In this section, we apply the proposed analytical framework to provide insights into how structure-function relationships evolve with age using the NKI dataset that comprises 196 healthy participants aged from 4 years to 85 years.

As the proposed SC-FC mapping is constructed via the functional mode with the largest eigenvalue, we first assess whether and how its contribution to the FC network varies with age. As shown in Fig. 5a, we find a weak but statistically significant increase in the contribution of this largest functional mode (r=0.16, FDR-corrected p=0.04), suggesting that the pattern of interregional functional interactions is increasingly governed by this inherent mode with age. As the first three functional modes almost explained the variance in empirical FC networks (Supplementary Fig. S1B; *R* = 0.91 *±* 0.03), we further assess age-related changes in the contributions of functional modes with the second and third largest eigenvalues. As shown in Fig. 5b, we find that none of them increase across the lifespan (2nd mode, r=-0.07, FDR-corrected p=0.36; 3rd mode, r=-0.16, FDR-corrected p=0.04). We also calculate the functional diversity (FD) for each participant, which measures the dispersion of the contribution of different functional modes (see Materials and Methods). Larger functional diversity indicates that the pattern of functional interactions is governed to a greater extent by distinct functional modes, whereby the distribution of functional modes’ contributions is closer to a uniform distribution. As shown in Fig. 5c, we observed a statistically significant association between FD and age (r=-0.15, p=0.03), indicating age-related decreases in functional diversity across the human lifespan. Taken together, these findings suggest that the diversity of functional modes gradually decreases with age, with the pattern of brain functional interactions increasingly dominated by the most contributing functional mode.

**Fig. 5.**
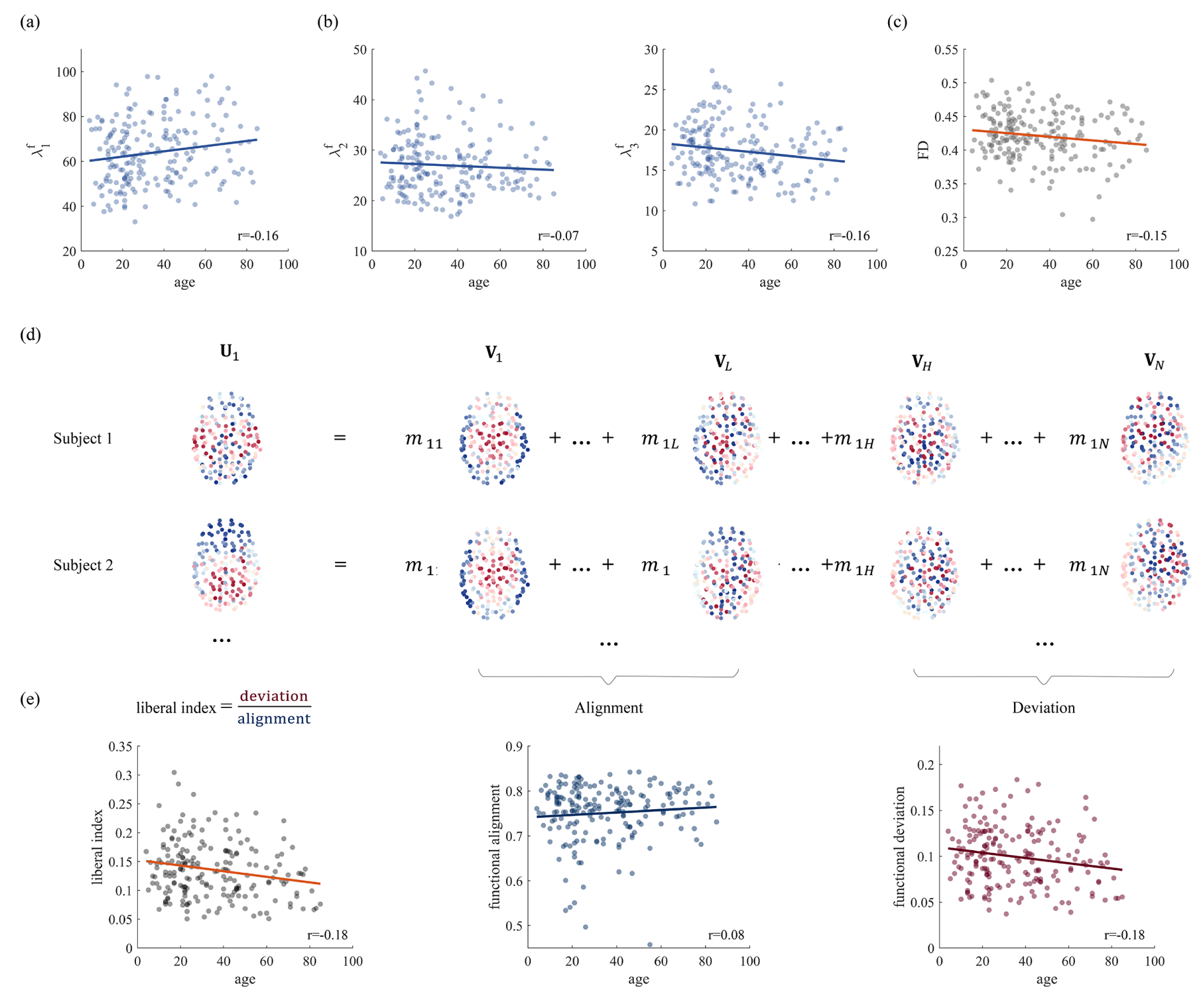
Age-related variations in SC-FC relationships within the proposed analytical framework. (**a**) The contribution of the first functional mode to the FC network increases with age. (**b**) Age-related variations in contributions of the 2nd or 3rd functional modes. (**c**) Decreases in functional diversity across the lifespan. (**d**) The functional interaction pattern for each subject is decomposed into structure-aligned and structure-deviated components using the eigenspectrum of the SC network. The ratio between the norms of these two components is used to measure structure-function liberality. (**e**) Reduced structure-function liberality across the lifespan, which is predominantly driven by the weakened functional deviation.

We next examine how structure-function coupling relationships vary with age within the proposed mapping framework. We utilize the structural eigenspectrum to decompose the functional mode that captures the essence of the FC network into two separate components: one is structure-aligned functional component, which potentially represents direct dependence on physical connections, and the other is structure-deviated component, which potentially reflects intermediate polysynaptic interactions in the structural network. (Fig. 5d; Materials and Methods). The norms of these two components quantify the extent to which functional interactions are organized in an aligned or deviated manner atop the underlying structural connectome. To investigate whether structure and function evolve synergistically or divergently throughout the human lifespan, we introduce a liberal index, which is estimated as the energy ratio of functional components decoupled (deviation) versus coupled (alignment) with structure, and correlate the values with the subjects’ ages. As shown in Fig. 5e, we observe a decline in the liberal index (*r* = *−*0.18*, p* = 0.01), suggesting that the liberality between brain structure and function gradually decreases with age. We also associate the magnitudes of functional alignment and functional deviation with subjects’ ages and find that age-related alterations in structure-function coupling relationships can be dissociable. We find that the magnitude of functional deviation decreases throughout the lifespan (*r* = *−*0.18*, p* = 0.01) whereas variability in functional alignment does not exhibit a statistically significant correlation with age (*r* = 0.08*, p* = 0.39). This indicates that even though both types of coupling patterns contribute to functional interactions, structure-deviated portions are the ones that reflect inter-individual variation during human brain development and aging.

## Discussion

Accurate quantification of structure-function relationships is imperative for understanding how cognitive functions emerge from the anatomical substrate and how the interrelationship of structural and functional connectivity varies across individuals with different traits and phenotypes. However, the connectome-based correspondence between structure and function is relatively moderate, despite a variety of statistical models (*8, 49*), communication models (*11, 12*), and biophysical models (*9, 10*). Emerging evidence suggests that biophysical models enriched with regional heterogeneity can be better fitted to brain functional connectivity (*27, 50*). Indeed, significantly higher prediction accuracies than previously implied have been achieved by the machine learning approach (*37*) which focuses on optimizing its performance rather than providing any mechanistic insights. However, these studies have eschewed simple models in favor of complex ones, and as a result, SC-FC mappings are constructed through time-consuming simulations or training procedures, with the essential principles underlying structure-function coupling remaining elusive.

Here, inspired by the graph spectrum analyses, we propose a novel mapping method that allows SC and FC networks to be tightly linked in a simple manner. Whereas most previous studies estimate the structure-function relationship based on putative communication mechanisms and neurophysiological processes, we simply exploit network constituent eigenmodes to enhance our understanding of how neuronal coactivation patterns emerge from indirect, collective interactions among brain regions. Using the eigendecomposition of the FC network, we first show that the essence of observed FC networks can be characterized by a just few inherent modes, i.e., those that possess large eigenvalues. This observation demonstrates the utility of eigenmodes for dimensionality reduction, which is key to establishing the proposed mapping framework. Typically, the functional mode with the largest eigenvalue is of particular interest in terms of its essential role in governing the formation of functional connectivity. By projecting this functional mode into the parsimonious basis formed by mutually orthogonal structural eigenmodes, we establish a concise and strong link between brain structural and functional networks, whose prediction performance is verified on two independent datasets (LAU and NKI).

Analogous to the recent machine learning approach (*37*), our study suggests that brain structure and function are indeed inextricably linked, with the predictability of FC significantly improved compared to those previously suggested by conventional eigenmode (*15*) and communication (*26*) methods at both whole-brain and regional levels. The key property that makes the proposed mapping stand out may be that our approach aims to capture the inherent patterns of functional interactions rather than to accurately fit functional connectivity between each pair of brain regions. This change yields a simple mapping procedure, where model parameters are solved in a completely analytical manner, and potentially suppresses the interference of spurious connections benefiting from the utilization of eigenmodes (*39*). Of note, although the conventional eigenmode approach also uses structural eigenmodes as input, the proposed method effectively embodies the collective, high-order properties of structure-function relationships through the introduction of cross-terms that describes the interaction, which appears to serve as the main contributors to the enhanced interpretation of FC networks. Intriguingly, regional heterogeneity is preserved in this enhanced structure-function correspondence, exhibiting a good agreement with the previous findings (*30, 31*) that structure and function are tightly coupled in primary sensory regions but diverge in polysensory association regions. This observation raises the possibility that regionally heterogeneous structure-function relationships may be an inherent brain property induced by hierarchical microscale organization, including cytoarchitecture (*51*), intracortical myelination (*34*), and laminar differentiation (*35*). One prominent account posits that the rapid evolutionary expansion of cortical mantle effectively releases association areas from early sensory-motor hierarchies, resulting in great signal variance and weak structure-function relationship in transmodal association cortex (*52*).

Importantly, we demonstrate that the proposed SC-FC mapping is sufficiently accurate to capture subject-specific information not explained by the mean in a less homogenous population, opening the possibility of a detailed investigation of individual differences in structure-function relationships. Consider, for instance, the effects of development and aging. Prior studies have reported age-related alterations in brain structural and functional connectomes, which are considered to be associated with cognitive performance during development and senescence (*53–56*). However, less is known about how structural and functional brain networks evolve jointly, especially from the perspective of their inherent constituent patterns. The proposed method provides an analytical framework in which we could explain the patterns of interregional functional interactions in terms of distinct structure-informed modes. Here, we focus on two typical functional components: one is constituted by structural modes with large positive eigenvalues and organized in close relation to structural connectivity, underscoring the direct dependence on anatomy; the other is represented by structural modes with negative eigenvalues and greatly untethered from the structural constraint, emphasizing the roles of indirect, high-order communication. By distinguishing the cases of ‘intermediate polysynaptic interactions’ (deviation) versus simple ‘signaling along the physical connections’ (alignment), we identify the age-sensitive components of functional connectivity. We find that structure-deviated functional components weaken with age whereas the magnitude of structure-aligned components is preserved with age, manifesting as gradually reduced structure-function liberality across the human lifespan. This observation is particularly interesting in the context of the prior finding that key information for individual identification is found in the functional component deviated from structure (*57*), implying that structure-function liberality may be an individually variable feature reflecting the inner workings of the brain. Furthermore, several previous studies have demonstrated correlations between the structure-function coupling relationship and inter-individual variability in cognitive traits (*18, 58–60*). Stronger structure-function coupling is found to be associated with better abilities of complex cognition, such as reasoning and cognitive switching, which may benefit from reliable and efficient information transmission (*18, 61*). In contrast, other cognitive traits, such as the level of awareness and attention maintenance, are considered to benefit from less alignment between structure and function, a configuration that might be instrumental for information integration across the brain (*23, 57*). Combined with these complementary roles of different degrees of structure-function alignment, our findings of age-related decreases in structure-deviated functional components may promote mechanistic insights concerning cognitive changes across the human lifespan. Our mapping framework also provides analytical tools to detect potentially behavior-sensitive or stimulus-sensitive components of functional interaction patterns, with significant implications in inferring physiological processes of cognitive functions and informing the diagnosis and treatment of psychiatric disorders.

There are several limitations and possible developments in this study. For example, while a concise and strong structure-function correspondence is established through the most contributing functional eigenmode, other functional modes (e.g., modes with 2nd and 3rd largest eigenvalues) also leave their signature on the formation of the FC network. Conceptually, future research could incorporate these functional modes into the proposed SC-FC mapping framework to obtain a more comprehensive understanding of how intricate functional connections emerge from the underlying structural network. In addition, our structure-function mapping does not embody the temporal fluctuations (*62*) or the regional heterogeneity (*50*) of the coupling patterns. Thus, another direction for future research is to enrich network constructions with dynamic interactions and microscale attributes, which would promote the richer interpretation of functional interactions among brain regions.

In conclusion, we demonstrate a novel SC-FC mapping method by which the patterns of functional interactions can be closely related to the organization of structural connections. The methodology, while feasible easily, provides informative quantification of individual-level relationships between structure and function, achieving substantially greater explanatory power for FC networks relative to previous mappings. It also offers a new avenue to understand how brain structural and functional networks are intertwined in terms of their constituent eigenmodes, holding great potential for an investigation of the evolving properties of the structure-function relationship across the lifespan. Alterations due to cognitive tasks, lesions, and neurological diseases might be another promising application of the proposed approach that would provide valuable insights.

## Materials and Methods

### Data

In this study, we performed all analyses in two independent datasets. The first one was collected by Department of Radiology, University Hospital Center and University of Lausanne (LAU) (*63*). This dataset included 70 healthy participants (27 females, 28.8±9.1 years old). Informed consent approved by the Ethics Committee of Clinical Research of the Faculty of Biology and Medicine, University of Lausanne was obtained from all participants. Diffusion spectrum images (DSI) were acquired on a 3-Tesla MRI scanner (Trio, Siemens Medical, Germany) using a 32-channel head-coil. The protocol was comprised of (1) a magnetization-prepared rapid acquisition gradient echo (MPRAGE) sequence sensitive to white/gray matter contrast (1-mm in-plane resolution, 1.2-mm slice thickness), (2) a DSI sequence (128 diffusion-weighted volumes and a single b0 volume, maximum b-value 8,000 s/mm2, 2.2×2.2×3.0 mm voxel size), and (3) a gradient echo EPI sequence sensitive to blood oxygen level-dependent (BOLD) contrast (3.3-mm in-plane resolution and slice thickness with a 0.3-mm gap, TR 1,920 ms, resulting in 280 images per participant). Gray matter was divided into 68 brain regions following Desikan–Killiany atlas (*64*) and further subdivided into 219 approximately equally sized nodes according to the Lausanne anatomical atlas using the method proposed by (*65*). Individual structural networks were constructed using deterministic streamline tractography, initiating 32 streamline propagations per diffusion direction for each white matter voxel (*66*). Functional networks were reconstructed using fMRI data from the same individuals. fMRI volumes were corrected for physiological variables, including regression of white matter, cerebrospinal fluid, and motion. fMRI time series were lowpass filtered. The first four volumes were discarded and motion “scrubbing” was performed (*67*). Functional connectivity matrices for individual participants were constructed by estimating the Pearson correlation between the fMRI time series of each pair of brain regions. More details regarding network construction can be obtained online at the LAU website (https://zenodo.org/record/2872624#.XOJqE99fhmM).

The second one was the Nathan Kline Institute (NKI)/Rockland Sample public dataset. This dataset consisted of 196 participants (82 females, age range=4-85). Informed consent approved by the Institutional Review Board was obtained from all participants (informed consent was also obtained from child participants and their legal guardians). The scan was performed in a Siemens Trio 3T scanner. The protocol consisted of: (1) 10-minute resting state fMRI scan (R-fMRI), (2) 6-direction diffusion tensor imaging (DTI) scan, (3) 64-direction diffusion tensor imaging scan (2mm isotropic), (4) MPRAGE anatomical scan, (5) MPRAGE anatomical scan SHORTER sequence, (6)T2 weighted sequence, (7) A variety of psychiatric, cognitive and behavioral assessments. The preprocessing contains head movement correction, denoising, and thresholding. The structural connectivity (SC) and functional connectivity networks, composed of 188 ROIs based on the Craddock 200 atlas (*68*), were derived from diffusion tensor imaging (DTI) and functional magnetic resonance imaging (fMRI), respectively. A comprehensive description of the data can be obtained online at the NKI website (http://fcon_1000.projects.nitrc.org/indi/pro/nki.html).

### Structural and functional modes

Applying an eigendecomposition, the FC network can be decomposed as **FC** = **UΛ***^f^* **U***^T^* where the eigenvalues are represented by 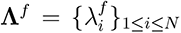 and eigenvectors are represented by **U** = **U_i_***i≤i≤N*. N indicates the number of network nodes. Several negative eigenvalues that may be induced by the noise were set to 0, which did not result in significant losses of information on FC networks (*69*). According to the spectral graph theory (*40*), these mutually orthogonal eigenvectors **U** can be interpreted as the N inherent constituent modes of the FC network. As all eigenmodes are scaled to the unit norm, the magnitude of eigenvalues mirrors the contribution of the corresponding functional mode to the FC network. Typically, the eigenmode with the largest eigenvalue represents the functional pattern that has the greatest impact on the formation of functional connectivity. Conversely, eigenmodes with zero eigenvalues are considered to have no contribution to FC, i.e. the FC network does not possess that functional mode. The alternative expression is 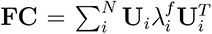, representing the FC network as the linear superposition of N independent functional modes. Similarly, the SC network can be decomposed as **SC** = **VΛ***^s^***V***^T^*, with the eigenvalues 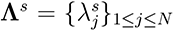 quantifying the smoothness (alignment) of the inherent modes specified by eigenvectors **V** = *{***V***_j_}*_1_*_≤j≤N_* (*18*). The 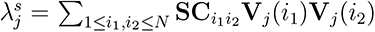 will be positive if the structural mode **V***_j_* is aligned to the underlying connectivity (values of most connected nodes possess same signs) and will be negative if the mode **V***_j_* deviates from the connectivity (values of most connected nodes possess different signs). We sorted these structural eigenvectors in descending order of eigen-values, constructing an eigenspectrum spanning from structural modes closely aligned with SC to modes deviated from SC.

### SC-FC mapping

The link between brain structural and functional networks can be constructed by projecting functional modes *{***U***_i_}*_1_*_≤i≤N_* into structural modes **V** = *{***V***_j_}*_1_*_≤j≤N_* :

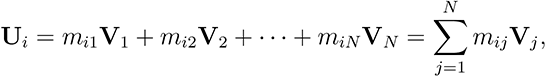

where parameters *{m_ij_}*_1_*_≤i,j≤N_* can be computed as 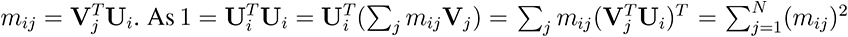, the magnitude of *m*^2^_ij_ can be used to reflect the contribution of structural mode **V***_j_* to functional mode **U***_i_* (*69*).

The FC network can thus be represented as:

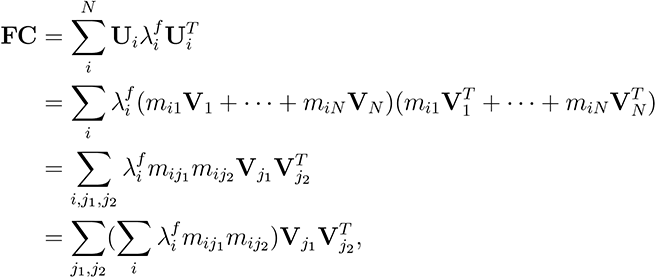

where 1 *≤ i, j*_1_*, j*_2_ *≤ N* . To avoid the overfitting issue, we only keep the functional mode with the largest eigenvalue given its essential role in governing the formation of functional connectivity. The estimation of the FC network is then simplified as:

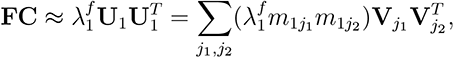

with the number of parameters is restricted to N. The parameters *{m*_1_*_j_}*_1_*_≤j≤N_* were estimated as 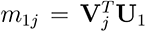. This expression can be considered as a generalization of the conventional eigenmode approach where the FC network is expressed as a weighted combination of structural eigenvectors 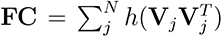. *h*(*·*) represents a linear function. The important difference is that the present approach introduces the interactions between different structural modes, i.e. the cross terms 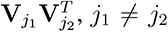. Furthermore, more refined quantification of the relationship between structure and function can be obtained by incorporating more functional modes into this analytical framework.

### Functional diversity

To measure the diversity of contributions of different functional modes, here called functional diversity (FD), we estimated the similarity of the empirical distribution of functional eigenvalues to the uniform distribution:

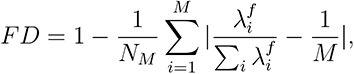

where M is the number of functional modes that the FC network possesses and *N_M_* = 2(*M −* 1)*/M* is a normalization factor that restricts the FD to the interval [0,1]. At one extreme where the FD value equals 0, the FC network is completely governed by one inherent mode; at the other extreme where the FD=1, all functional modes contribute equally to the formation of functional interactions.

### Structure-function liberality

Within the present analytical framework, the inherent pattern of functional interactions can be investigated in the context of a structural eigenspectrum spanning from modes closely aligned to anatomical connections (those with positive structural eigenvalues) to modes deviated from the anatomy (those with negative structural eigenvalues). Here, we exploite the Graph Fourier Transform (GFT) (*40*) and spectral filtering to split the most contributing functional mode into two separate components: one represented by the first *L_A_* structural modes, exhibiting tight coupling with the structure, and the other represented by the last *L_D_* structural modes, exhibiting flexible deviations from the structural substrate. That is,

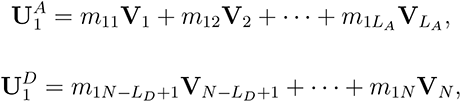

where **U**_1_^A^ and **U**_1_^D^ denote structure-aligned and structure-deviated components of the functional mode, respectively. *{m*_1_*_j_}*_1_*_≤j≤N_* are parameters estimated in SC-FC mapping procedure. Considering that there is no general method to determine the threshold *L_A_* and *L_D_*, we chose a default value (*L_A_*=*L_D_* =10) following the previous literature (*18*), and performed a sensitivity analysis to confirm the robustness of results to threshold selection (Supplementary Fig. S5). The intensity of the aligned and deviated portions was measured as the norms of **U***_1_^A^* and **U***_1_^D^* . We further introduce the structure-function liberality, which is estimated by the energy ratio between the structure-aligned and structure-deviated components, to identify to what degree the functional interaction pattern is misaligned versus aligned with the structure. Correlating this liberal index with the age, we could explore the evolving property of structure-function relationships across the human lifespan. There existed two distinct possibilities: (1) the structure-function liberality was preserved with age; (2) the structure-function liberality exhibited age-related change. The first one indicates that the structure-function relationship is preserved with age, implying that lifespan differences in FC networks may be simply induced by changes in structural architecture. The second one suggests that SC and FC networks change divergently with age, with increasing or decreasing liberality indicating that functional interaction patterns are gradually untethered or tethered by structural constraints.

## Acknowledgments

This work is supported by National Key Research and Development Program of China Grant No. 2022AAA010030, 2021YFB2700304, and Program of National Natural Science Foundation of China Grant No. 62141605, 12201026, 11922102, 11871004.

## Author contributions

Conceptualization: Y.Y., X.W.

Methodology: Y.Y., Z.Z., L.L., X.W.

Investigation: Y.Y., Z.Z., L.L., H.Z., Y.Z., Y.Z., X.W., S.T.

Visualization: Y.Y., Y.Z.

Supervision: X.W., S.T.

Writing—original draft: Y.Y.

Writing—review and editing: X.W., S.T.

## Data and materials availability

The Lausanne dataset is available at https://zenodo.org/record/2872624#.XOJqE99fhmM. The Nathan Kline Institute (NKI)/Rockland Sample public dataset is available at http://fcon_1000.projects.nitrc.org/indi/pro/nki.html.

---

**Fig. S1.**
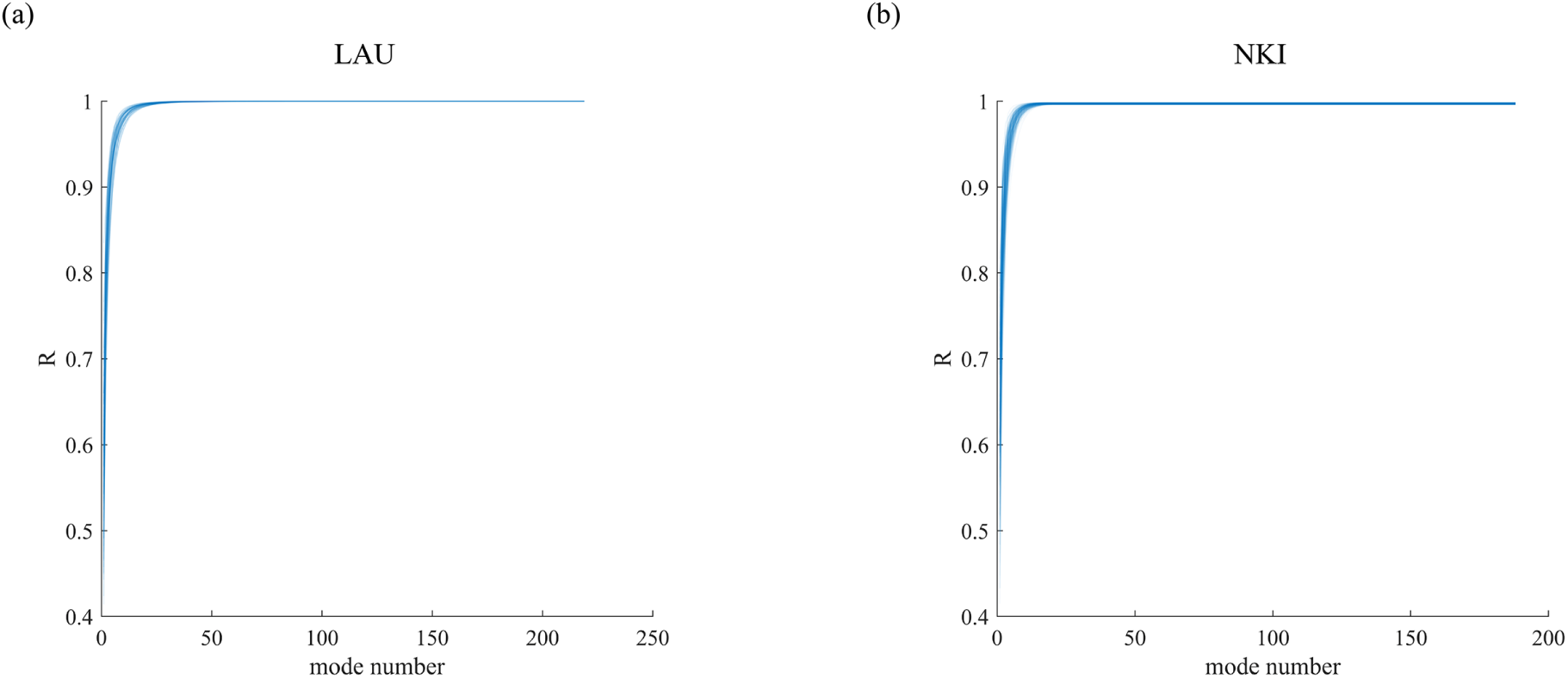
Prediction accuracy R as a function of the number of functional eigenmodes. (**a**) R for all subjects in LAU dataset. (**b**) R for all subjects in NKI dataset. Note that the prediction accuracy increases dramatically with the number of functional eigenmodes exploited in the proposed mapping, and that the first three functional modes almost explain the variance in the empirical FC networks (*R* = 0.86 *±* 0.04 in LAU and *R* = 0.91 *±* 0.03 in NKI).

**Fig. S2.**
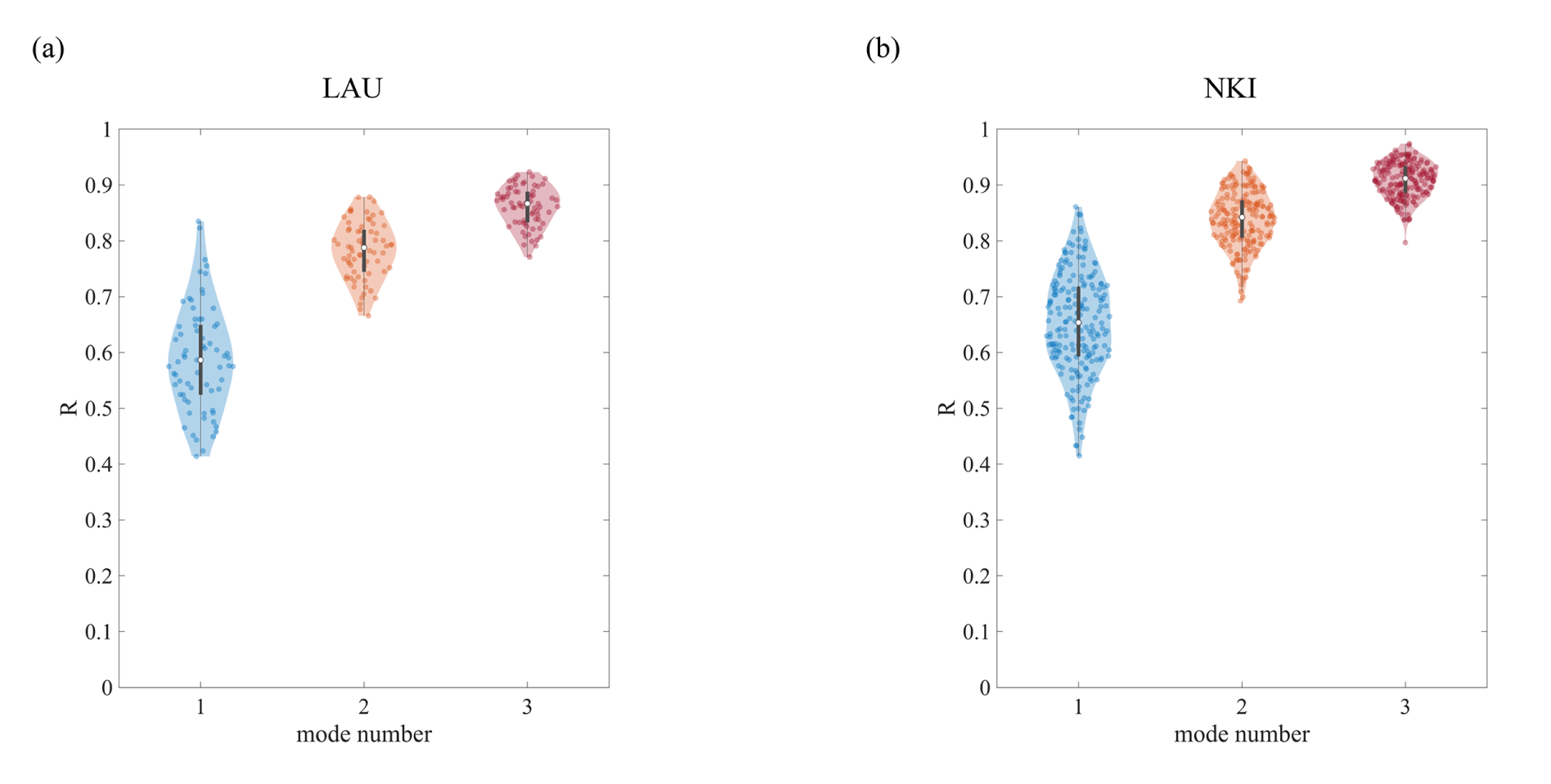
The prediction accuracy of the proposed mapping incorporating different numbers of functional modes. In each violin plot, the box indicates the interquartile range and the empty circle indicates the median value. Points represent individual subjects. (**a**) Prediction accuracy R for all subjects in LAU dataset. Specifically, *R* = 0.59 *±* 0.09 when incorporating the first one functional modes; *R* = 0.78 *±* 0.05 for the first two functional modes; *R* = 0.86 *±* 0.03 for the first three functional modes. (**b**) Prediction accuracy R for all subjects in NKI dataset. Specifically, *R* = 0.65 *±* 0.09 when incorporating the first one functional modes; *R* = 0.84 *±* 0.05 for the first two functional modes; *R* = 0.91 *±* 0.03 for the first three functional modes.

**Fig. S3.**
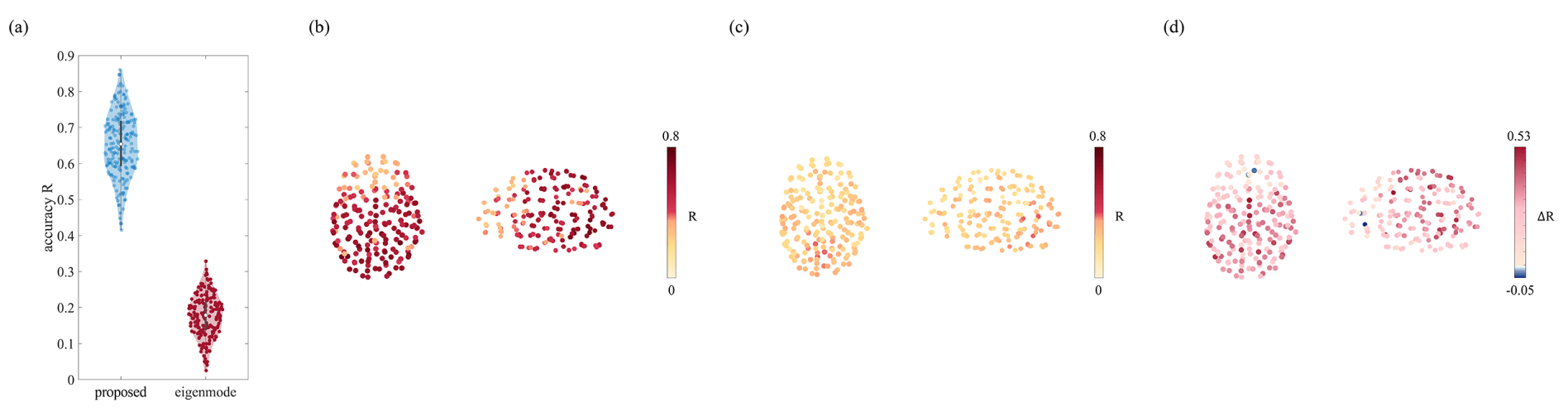
The performance of the proposed model in NKI dataset. (**a**) The prediction accuracy of the proposed method vs. the conventional eigenmode approach for all individual subjects. In each violin plot, the box indicates the interquartile range and the empty circle indicates the median value. (**b**) The spatial pattern of SC-FC coupling estimated by the proposed method; *R* = 0.65 *±* 0.09. (**c**) The spatial pattern of SC-FC coupling estimated by the eigen-mode approach; *R* = 0.18 *±* 0.06. (**d**) Regional differences between prediction accuracies of the proposed and eigenmode methods. 77 *±* 8% of brain regions are better explained by the proposed mapping.

**Fig. S4.**
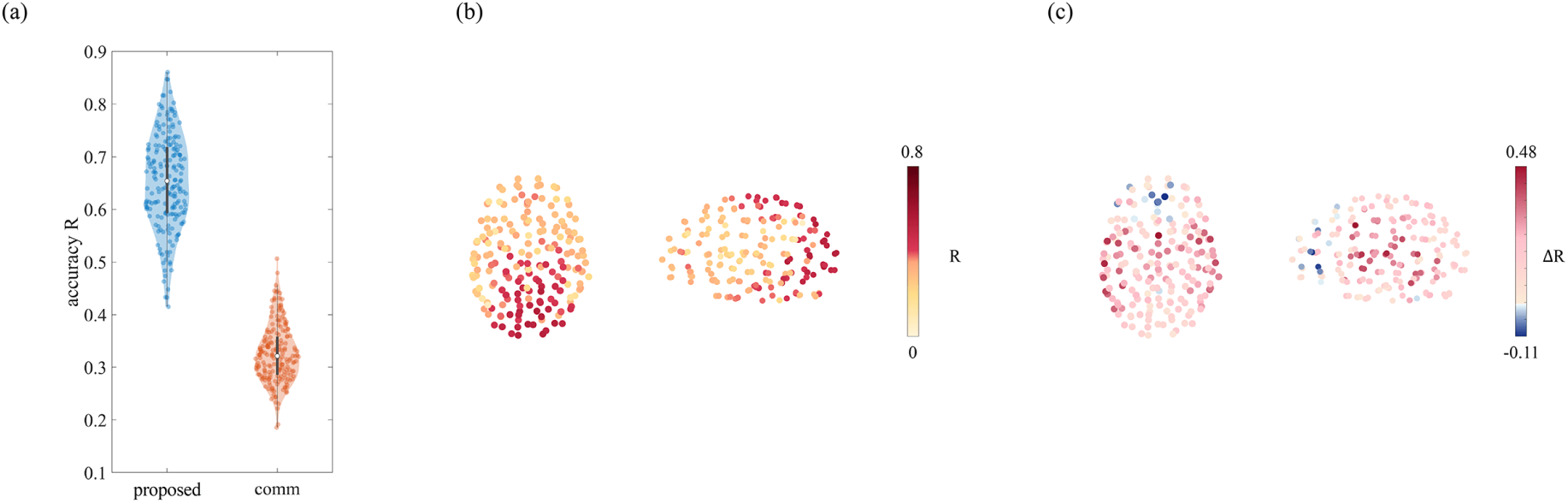
Comparison with the communication model in NKI dataset. (**a**) The prediction accuracy of the proposed method vs. the communication model for all individual subjects. In each violin plot, the box indicates the interquartile range and the empty circle indicates the median value. (**b**) The spatial pattern of SC-FC coupling estimated by the communication model; *R* = 0.33*±*0.05. (**c**) Regional differences between prediction accuracies of the proposed and communication methods. 70 *±* 10% of brain regions are better explained by the proposed mapping.

**Fig. S5.**
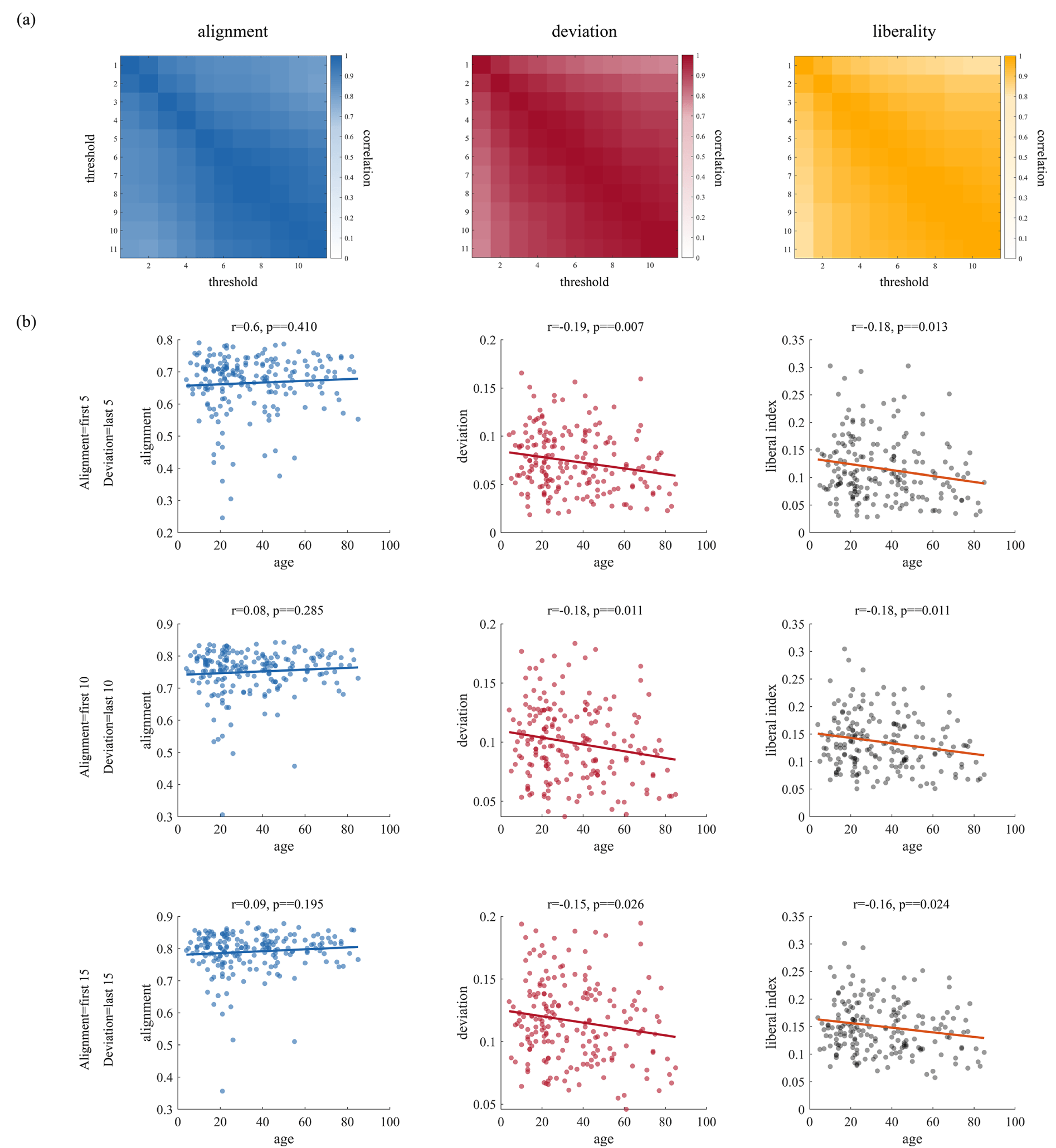
Sensitivity analysis of threshold selection. The results in the main text are reported with the threshold *K_L_* and *K_H_* equal to 10, that is, we use the first 10 structural eigenmodes to represent anatomy-aligned functional components and the last 10 structural eigenmodes to represent anatomy-deviated functional components. To test the robustness of the results, we repeat analyses under values of *K_L_* and *K_H_* from five below to five above the default values. (**a**) The correlation matrices among functional alignment (left panel), functional deviation (middle panel), and structure-function liberality (right panel) across different thresholds. We find these measures exhibit high stability, with the correlation coefficient *r* = 0.92 *±* 0.06 for functional deviation, *r* = 0.93 *±* 0.06 for functional alignment, and *r* = 0.94 *±* 0.05 for structure-function liberality. (**b**) Age-related variations in functional alignment (left panel), functional deviation (middle panel), and structure-function liberality (right panel) across the lifespan. The observations are robust to different choices of thresholds.

